# Protocol-dependence of middle cerebral artery dilation to modest hypercapnia

**DOI:** 10.1101/2021.03.11.434655

**Authors:** Baraa K. Al-Khazraji, Sagar Buch, Mason Kadem, Brad J. Matushewski, Kambiz Norozi, Ravi S. Menon, J. Kevin Shoemaker

**Author notes:** ***Correspondence*** J. Kevin Shoemaker, PhD, The University of Western Ontario, London, Ontario, Canada, N6A 5K7.

## Abstract

There is a need for improved understanding of how different cerebrovascular reactivity (CVR) protocols affect vascular cross-sectional area (CSA) when measures of vascular CSA are not feasible. In human participants, we delivered ~±4mmHg end-tidal partial pressure of CO_2_ (PETCO_2_) relative to baseline through controlled delivery, and measured changes in middle cerebral artery (MCA) cross-sectional area (CSA; magnetic resonance imaging (7 Tesla MRI)), blood velocity (transcranial Doppler and Phase contrast MRI), and calculated CVR based on steady-state versus a ramp protocol during two protocols: a 3-minute steady-state (+4mmHg PETCO_2_) and a ramp (delta of −3 to +4mmHg of PETCO_2_). We observed that 1) the MCA did not dilate during the ramp protocol, but did dilate during steady-state hypercapnia, and 2) MCA blood velocity CVR was similar between ramp and steady-state hypercapnia protocols, although calculated MCA blood flow CVR was greater during steady-state hypercapnia than during ramp, the discrepancy due to MCA CSA changes during steady-state hypercapnia. Due to the ability to achieve similar levels of MCA blood velocity CVR as steady-state hypercapnia, the lack of change in MCA cross-sectional area, and the minimal expected change in blood pressure, we propose that a ramp model, across a delta of ~−3 to +4mmHg PETCO_2_, may provide one alternative approach to collecting CVR measures in young adults with TCD when CSA measures are not feasible.

## Introduction

Cerebrovascular reactivity (CVR) studies assess changes in cerebral blood flow to a known vasoreactive stimulus (e.g., changes in end-tidal partial pressure of CO_2_; PETCO_2_). Measures of CVR are important because attenuated CVR may reflect preclinical vascular pathophysiology and an increased risk of mortality independent from cardiovascular risk factors or stroke incidence (1). The most commonly used technique for CVR measures in humans, transcranial Doppler (TCD) ultrasonography, provides an index of vascular blood flow changes (i.e., blood velocity) because the vascular cross-sectional area (CSA) values required for blood flow calculations (i.e., the product of CSA and blood velocity) are not collected with TCD. Thus, an assumption of an unchanging CSA is typically accepted, raising concern if changes in CSA do occur (2). To circumvent issues related to TCD measures of CVR, some research groups measure all four brain-supply (i.e., carotid and vertebral) arteries outside of the brain (3), or use expensive neuroimaging approaches (4,5).

An additional concern regarding quantification of CVR is the potential for changes in central hemodynamics during hypercapnia (elevated PETCO_2_) that could elevate cerebral blood flow due to changes in cardiac output (6) and blood pressure (6,7) and not directly due to cerebrovascular dilation (7). Another complicating factor in quantifying CVR between groups is potential variation in large cerebral artery reactivity, particularly when comparing age differences (8). As an example, our group’s previous work showed that compared to younger adults, older adults exhibited attenuated changes in large cerebral artery CSA in response to steady-state hypercapnia (9). However, obtaining cerebral artery CSA data requires access to costly MRI or CT systems. We aim to understand protocol designs that provide accurate CVR estimates when using TCD methods. The “ideal” velocity based CVR protocols conducted using TCD would require: 1) minimal CSA changes by conducting CVR protocols that result in negligible change in CSA, and 2) minimal influence of confounding variables such as blood pressure.

In the current study, we tested the hypothesis that a ramp (i.e., linear) CVR protocol within the ±5 mmHg range of relative changes in PETCO_2_ would provide minimal changes in CSA while still replicating CVR outcomes from the more standard steady-state hypercapnia CVR protocol. Our rationale for this range of relative changes in PETCO_2_ comes from the emerging knowledge of a sigmoidal change in MCA CSA, with minimal changes in hypercapnia, within the −5 to +5 mmHg from resting PETCO_2_ (10). We chose a ramp-style CVR protocol to compare to CVR measures from a steady-state hypercapnia protocol (0 to ~5 mmHg) because: 1) the ramp protocol is a well-established protocol for CVR measures (7,11,12), and 2) existing means for calculating CVR from standard steady-state protocols use linear slope methods which simply reduce to a ramp design. We acknowledge that the ramp protocol within the ~Δ±5 mmHg from baseline PETCO_2_ range may potentially affect vascular dilation differently than a steady-state hypercapnia protocol as the hypocapnia portion preceding hypercapnia may blunt blood velocity CVR (13). A sub-analysis in our earlier work, however, indicated that order of condition did not affect CSA reactivity in young adults (4). Additionally, existing methods primarily focus on velocity-based CVR without considering CSA, and we wanted to design a CVR protocol that minimized CSA changes (even if it involved hypocapnia) and blood pressure changes while retaining its ability to elevate cerebral blood flow (i.e., changing blood velocity). Other models can be considered but we want to test the ramp protocol as one example of alternative CVR designs that may elicit negligible CSA and BP changes when measuring blood velocity CVR using TCD.

To achieve the high temporal resolution of the MCA CSA for the required study, we developed a dynamic anatomical imaging sequence with high-temporal and spatial resolution to capture MCA CSA changes every 14 seconds throughout each vasoreactivity protocol (~±Δ4 mmHg) using 7T magnetic resonance imaging. Our objectives were to assess whether between steady-state and ramp protocols: 1) the MCA CSA increased compared to baseline, 2) blood pressure remained stable throughout duration of the protocol, and 3) MCA blood velocity (via TCD alone), and calculated flow vasoreactivity were similar.

## Materials and Methods

### Participants

All testing was conducted at the Centre for Functional and Metabolic Mapping at The University of Western Ontario. The Human Subjects Research Ethics Board at the University of Western Ontario (London, Ontario, Canada) approved the experiment protocols herein. Informed consent from 12 healthy subjects (19-25 years of age; 6 males) was obtained prior to scanning. A sample size calculation was based off of our previous work with blood velocity reactivity measured with TCD during steady-state hypercapnia (4). Specifically, for a within-subject design, with a Cohen’s *d* of 1.02, alpha level of significance of 0.05, and statistical power of 0.80, we calculated a sample size of 10 and recruited 12 individuals due to our laboratory’s expected attrition rate of 10-12% with our neuroimaging studies. Participants were ineligible if they were smokers, pregnant, or had any of the following conditions: Raynaud’s disease, respiratory illnesses, diabetes, claustrophobia, history of psychosis, eating disorders, manic or bipolar disorder, major psychiatric conditions, or dependence on alcohol or drugs.

### Procedure and data recording

Testing was completed between 10am – 2pm. Participants refrained from exercise, alcohol, drugs, and caffeine within 12 hours prior to testing. We used TCD and MRI to assess the cerebral vasoreactivity in response to steady-state (three minutes) bouts of hypercapnia (HC), and a ramp protocol from hypocapnia to hypercapnia (four minutes) (Fig. 1).

**Figure 1.**
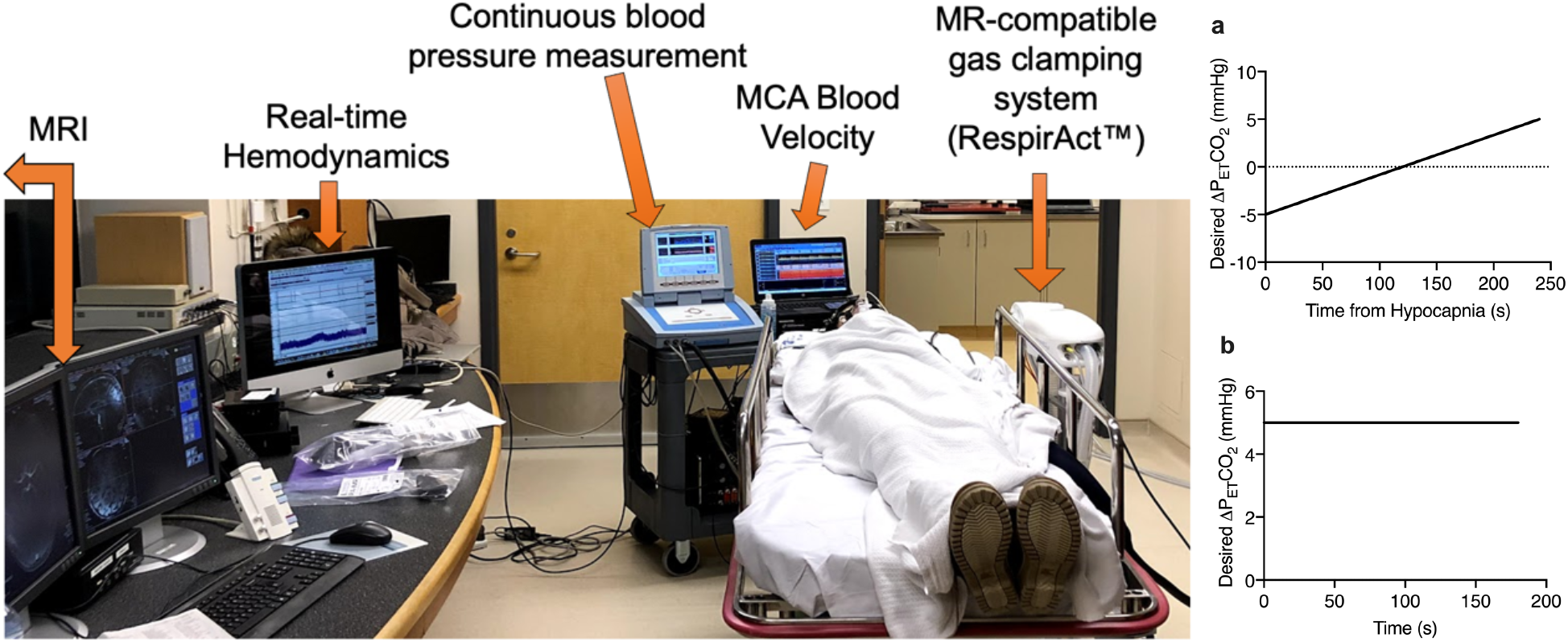
Experimental protocol schematic. Left: Experimental setup displaying real-time data collection for both transcranial Doppler and MRI sessions. Right, panel a: Ramp hypercapnia protocol with a desired target range of −5 to +5 mmHg from baseline PETCO_2_ and a duration of 240 seconds. Right, panel b: Steady-state hypercapnia (SSHC) protocol with a target range of +5 mmHg from baseline PETCO_2_ and a duration of 180 seconds. Protocols were executed using the RespirAct™ device with preset protocols programmed with the desired PETCO_2_ values.

Steady-state and ramp protocols were each conducted twice, once for each of the TCD and MRI portions of the testing sessions and the order of TCD and MRI trials was randomized across participants. Unfortunately, we were unable to randomize the order of CVR protocols as it was difficult to stop and restart the RespirAct™ (Thornhill Research Inc., Toronto, Ontario, Canada) without doing extensive recalibration. The desired ventilatory rate was set to 12 breaths/min using a visual metronome for each session and was projected on a screen during the MRI scan. Our goal was to have the protocols fall within the ±5 mmHg from baseline PETCO_2_ range. Following a familiarization period of four minutes, the order and duration of protocols occurred as follows: 1) baseline (1 minute), 2) steady-state hypercapnia (target was +5 mmHg, although only reached ~+4mmHg; three minutes), 3) recovery (2 minutes) 4) baseline (1 minute), 5) −5 mmHg PETCO_2_ hypocapnia (brief hyperventilation; target was +5mmHg, although only reached ~ −3mmHg) 30 seconds), 6) incremental increase (ramp) from ~ −3 mmHg hypocapnia to ~ +4 mmHg relative PETCO_2_ hypercapnia (four minutes), 7) recovery (two minutes).

#### Manipulating target PETCO_2_ stimulus

Prior to the MRI scan, participants were fitted with a facemask attached to the RespirAct™ system, a modified sequential gas delivery breathing circuit(13) was used to clamp PETCO_2_ levels at the desired +5 or −5 mmHg (depending on protocol). Breathing rate and tidal volumes were calibrated prior to starting the breathing sequence.

#### MCA blood velocity and systemic blood pressure

While supine, continuous beat-to-beat arterial blood pressure was monitored using a Finapres® Finometer system, where a finger cuff was placed on the middle phalange of the third finger, and the finger blood pressure was calibrated with an upper arm cuff (Finapres® Medical Systems, Amsterdam, Netherlands). The MCA was insonated with a 2 MHz ultrasound probe placed at the temporal window and the peak blood flow velocity envelope was collected using the Neurovision TCD System (Multigon Industries Inc., NY, USA). All analog data were sampled at 1000 Hz using the PowerLab data acquisition system (ADInstruments, Dunedin, Otago, New Zealand).

#### MCA vascular diameter and blood velocity during MRI

A 7 Tesla MRI (Siemens, Magnetom Step 2.3, Erlangen, Germany) system was used to acquire the following datasets: 1) 3D time-of-flight (TOF) with 0.8mm isotropic voxel resolution, echo time (TE)/ repetition time (TR) = 2.59ms/18ms, flip angle (FA) = 15°, bandwidth (BW) = 203Hz/pxl; 2) single-slice 2D phase contrast (PC-MRI) for MCA M1 segment blood velocity, with a voxel resolution of 0.3×0.3×1.4mm^3^, TE/TR = 7.72ms/24.3ms, four averages, FA = 20°, velocity-encoding (Venc) = 100cm/s and BW = 250Hz/pxl. During PC-MRI, a Venc value of 100cm/s was used for all subjects, except for one hypercapnic case, where a Venc of 130cm/s was used to avoid wrap-around artifact; and 3) cross-sectional area of the MCA M1 segment using single-slice 2D turbo spin-echo T1-weighted imaging with 0.5×0.5×1.5 mm^3^, TE/TR = 12/750ms, BW = 270Hz/pxl, with an acquisition time of 13-14 seconds. The TOF data were used to locate a straight segment on the right MCA M1 segment with the least curvature. The single-slice PC-MRI and T1-weighted data were then acquired orthogonally to the axis of the selected MCA segment. The T1-weighted data were acquired sequentially in order to monitor the changes in MCA diameter.

### Data analysis

Data analysis was carried out offline using custom R scripts (RStudio; v. 2020), GraphPad (V.8), and LabChart Pro (v.8, ADInstruments, Dunedin, Otago, New Zealand).

#### Transcranial Doppler ultrasound

The MCA blood velocity (via TCD) and MAP were averaged beat-by-beat over the cardiac cycle then were exported at 5 Hz sampling frequency and saved as text files (LabChart Pro v.8, ADInstruments, Dunedin, Otago, New Zealand). During the TCD collection phase of testing, start and end times of each event throughout the experimental breathing protocol were chronicled by comments added in the LabChart file.

#### Magnetic resonance imaging (MRI; 7 Tesla)

During the MRI collection phase of testing, start and end times of each event were chronicled based on the time associated with the desired PETCO_2_ on the exported RespirAct™ data file. For the T1-weighted anatomical MCA images, start and end times for each protocol event were recorded based off the MRI console such that images were lined up offline based on the DICOM image acquisition time. The brain anatomical images were imported into a DICOM reader, OsiriX software (Pixmeo©, Bernex, Switzerland), and MCA cross-sectional area (CSA) was measured by a blinded rater (MK) and compared against an expert rater (BKA).

The PC-MRI data were acquired for 30 seconds at baseline and within a ~70-second window following 50-60 seconds from the start of steady-state hypercapnia (as noted on Fig 4, panel J with the PC-MRI text bar). The MCA blood velocity measurements during the MRI session were obtained from PC-MRI data by manually contouring a region-of-interest (ROI) inside the MCA lumen. The contours were drawn using the software, signal processing in NMR (SPIN-Research, MR Innovations Inc., Detroit, MI, USA). Care was taken to avoid any peripheral voxels within the MCA lumen. The magnitude PC-MRI data was used to locate the MCA lumen. The peak velocities were calculated for each subject for baseline and hypercapnic states.

#### Data extraction

The LabChart text files, the TCD and MRI session RespirAct™ breath-by-breath PETCO_2_ values, and the measured CSA were aligned using RStudio (v. 2020) (14) for data extraction from specific epochs as indicated by the event comments in each file. The “print” function in the “magicfor” package in R (15) was used to extract data with each loop iteration for each participant, and variables of interest were exported as. csv files and imported to GraphPad Prism for graphing and analysis.

Steady-state condition time courses were baseline corrected by subtracting mean baseline value (−30 to 0 seconds of time window of interest) for each respective variable and plotted as delta values (Fig. 4). For the baselines and steady-state condition, the data were extracted from the following sections within the protocol: 1) 30 seconds baseline prior to onset of hypocapnia prior to the ramp protocol and 2) 30 seconds baseline prior to steady-state hypercapnia, and 3) 60 seconds at the end of steady-state hypercapnia. The ramp slope analysis included the entire hypocapnic to hypercapnic incremental data (see below).

#### Ramp protocol

The target PETCO_2_ and ramp protocol schematic is shown in Fig. 1, panel a. The target ventilation rate was 12 breaths/minute and participants were coached using a visual metronome. In order to equalize the spacing on the time axis when plotting the achieved PETCO_2_ data, the PETCO_2_ and corresponding time vectors were resampled to a fixed 12 breaths/minute sampling rate using the base R “approx” function in RStudio. Thus, two time vectors were created: 1) a target time vector that is based off of a 12 breath/minute ventilation rate and 2) a “fixed” time vector that is based off of resampling each participants data to meet the target time vector. The inter-individual differences for the “fixed” time vector are indicated by horizontal error bars (mean ± S.D.; Fig. 3, panels a-b), and the data are plotted and averaged for all participants along the target 12 breaths/minute time vector.

MCA blood velocity and systemic MAP data (in 0.2 second increments or 5 MHz sampling frequency) were plotted from the hypocapnic state to the hypercapnic state during the ramp protocol and averaged at each time point across the 12 participants (Fig. 3 panels c-d). Similarly, the CSA values along the ramp protocol were averaged for each time point across participants (Fig. 3 panel e). Although our goal was to collect 4 minutes of ramp data, the transition point from the nadir of hypocapnia to start of ramp was different for each person and it generally took approximately 2 breaths (~12 seconds) to sync with the desired PETCO_2_ for the hypercapnic ramp. Thus, to ensure the same number of samples (n=12) for each time point along the ramp protocol, we only extracted the last 228 seconds of ramp data for all participants (instead of the full 240 seconds).

#### Steady-state protocol

Data extraction and organization were similar to the ramp protocol except data extraction occurred between the start and end of the three-minute steady-state hypercapnia stimulus. As previously mentioned, the PC-MRI data acquisition commenced 50-60 seconds from start of steady-state hypercapnia. Thus, the continuous T1-anatomical MCA CSA scans were interrupted to allow for PC-MRI imaging (correlation with TCD measures of blood velocity are shown in Fig. 2). PC-MRI data were collected for 11 out of our 12 participants. As there were shifts in PC-MRI data collection start and end times across participants, the upper and lower bounds of these time points are indicated under the “PC-MRI” text bar on Fig. 3 (panel e, bottom row) to indicate to the reader that the mean and S.D. for CSA values in this portion of the protocol do not include all 11 participants (i.e., n < 11) for the CSA data.

**Figure 2.**
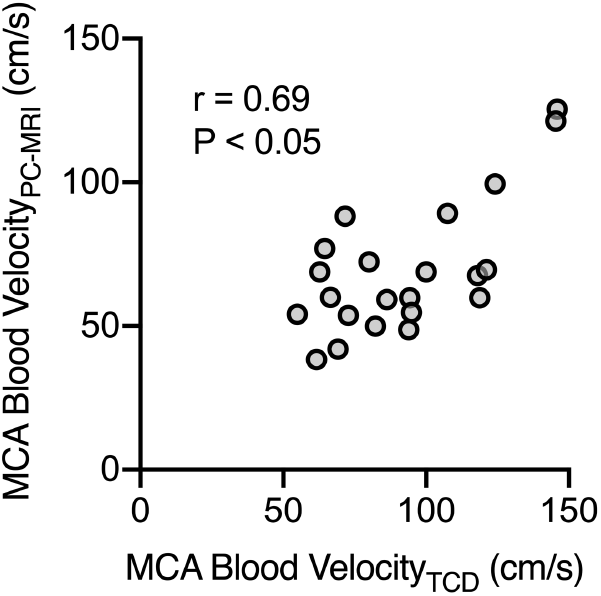
MCA blood velocity measure comparison between TCD and PC-MRI. Correlation plot comparing MCA blood velocity measured from transcranial Doppler (TCD) and phase contrast magnetic resonance imaging (PC-MRI). Baseline measures for both TCD and MRI sessions and steady-state hypercapnia measure at the ~1-2 minute mark for PC-MRI and in the last minute of steady-state hypercapnia for the TCD session for 22 pairs (n=11; r = 0.69, P<0.05).

**Figure 3.**
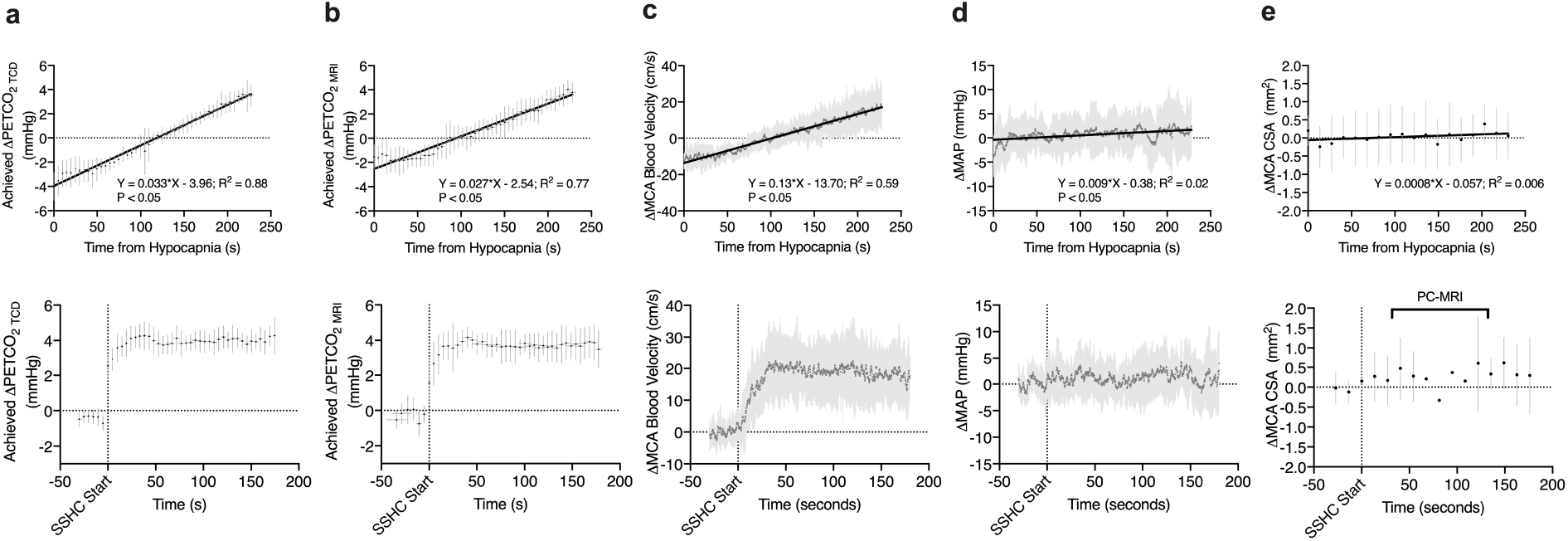
Measured variables during the ramp and steady-state hypercapnia protocols. Ramp protocol (228 seconds; top row) responses and steady-state hypercapnia (180 seconds; bottom row) for delta changes in each variable from baseline levels are shown. Panel a: achieved PETCO_2_ for TCD session, panel b: achieved PETCO_2_ for MRI session, panel c: MCA blood velocity in TCD session, panel d: mean arterial pressure (MAP), and panel e: MCA cross-sectional area (CSA). The “PC-MRI” text on panel e bottom row indicates the variable time window in which PC-MRI images were acquired and there was an interruption in the consistent MCA CSA measurements. A 30 second baseline is shown for the steady-state hypercapnia condition (bottom row) and hypercapnia is indicated by SSHC on x-axis with a vertical line. Linear regressions (panels a-e top row) are shown for each variable in the ramp protocol and a significant (non-zero; α level significance 0.05) slope is indicated by P < 0.5. N = 12 for all variables with data presented as mean±S.D.

The averaged raw values for the 30 seconds baseline (prior to start of stead-state hypercapnia; indicated as B on x-axis in Fig. 4) and last minute of steady-state hypercapnia (indicated as SSHC on x-axis on Fig. 4) are shown in Fig. 4 for PETCO_2_, MCA blood velocity (via TCD) and mean arterial pressure (MAP) during the TCD session and the PETCO_2_, MCA blood velocity (via PC-MRI) and MCA CSA during the MRI session.

**Figure 4.**
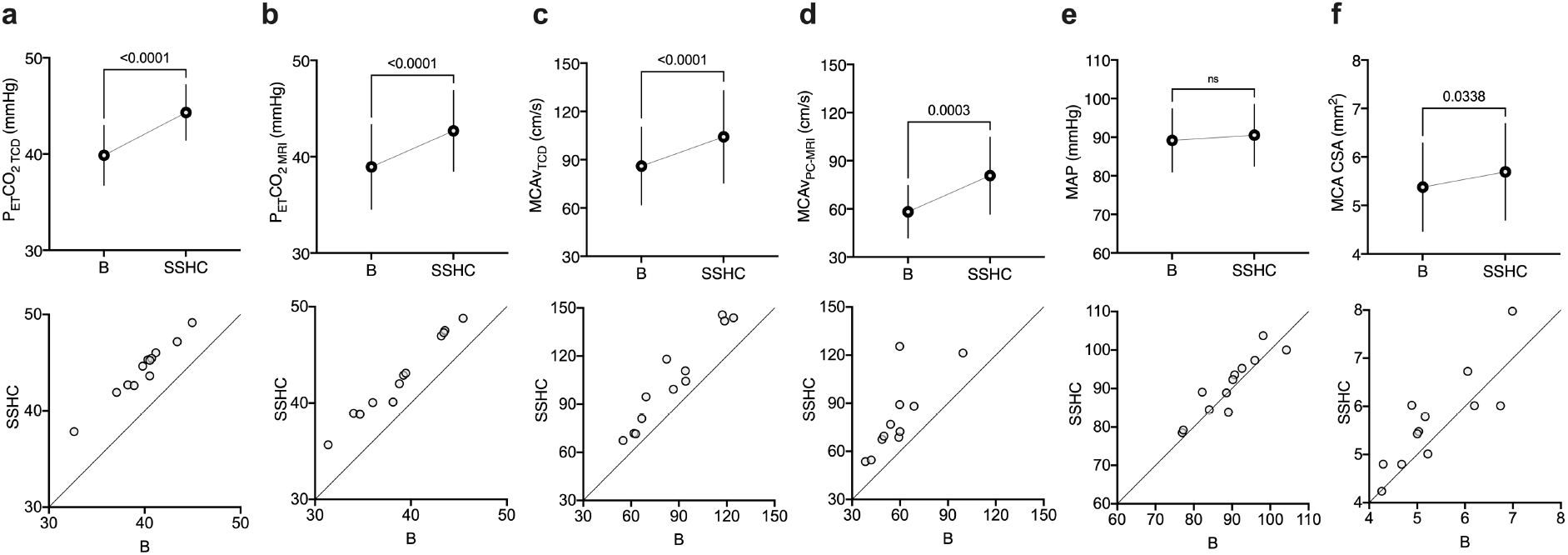
Baseline (B) and steady-state hypercapnia (SSHC) comparisons. Top row: Baseline (B) and steady-state hypercapnia (SSHC) measures (n=12 except for panel d where n=11, mean±S.D.). Where statistical significance occurs, p-values are indicated above data (ns = not significant; paired t-test; x03B1; level significance 0.05). Bottom row: Individual data points (n=12 except for panel d where n=11) comparing baseline (B) to steady-state hypercapnia (SSHC) responses for the same variable in that column as top row. The diagonal line (identity line) indicates where data would fall if there was no measurable effect of SSHC from B. Data above the identity line indicates an increase in variable measure with SSHC (from baseline; B).

#### MCA blood velocity reactivity calculations

The ramp and steady-state hypercapnia MCA blood velocity cerebrovascular reactivity (CVR) measures are shown in Fig. 5 panel a. For the ramp protocol, the MCA blood velocity slope (Fig.3 panel c) and the PETCO_2_ (for TCD session; Fig. 3 panel a) slope were calculated for each person and the MCA blood velocity CVR was calculated as:

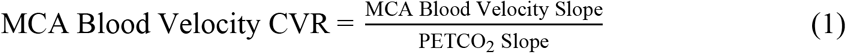

**Figure 5.**
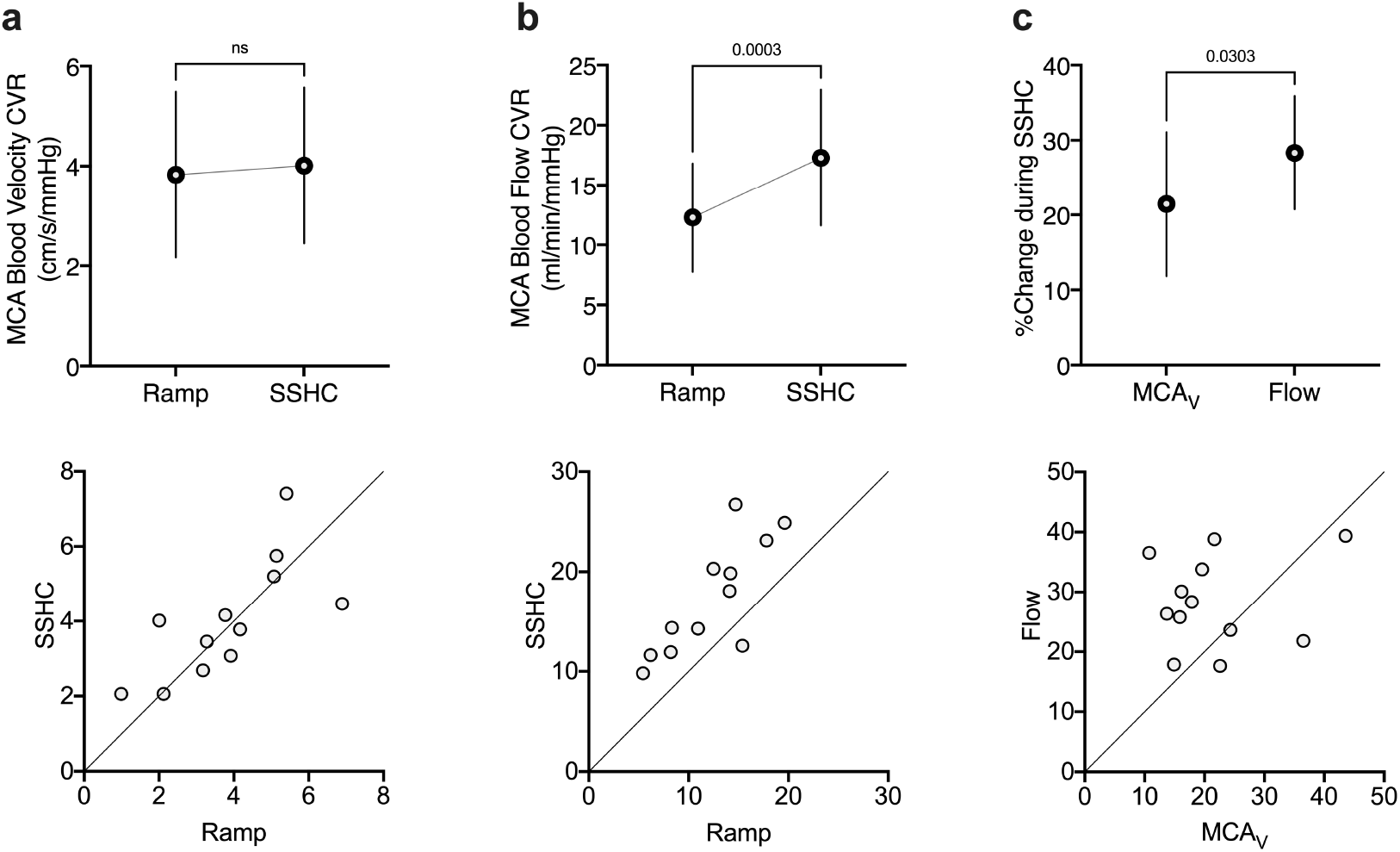
Middle cerebral artery (MCA) blood velocity and flow reactivity across protocols. Top row: Calculated MCA blood velocity cerebrovascular reactivity (CVR) was not different between ramp and steady-state hypercapnia (SSHC; panel a), while MCA blood flow CVR was higher in SSHC compared to the ramp protocol (panel b). The percent change in calculated MCA blood flow from baseline was higher than the percent change in MCA blood velocity from baseline (panel c) with SSHC. Where statistical significance occurs, p-values are indicated above data (ns = not significant; paired t-test; x03B1; level significance 0.05). N=12 for all variables with data presented as mean±S.D. Bottom row: Individual data points (n=12) comparing ramp to steady-state hypercapnia (SSHC) responses (panels a-b) for the same variable in that column as top row or MCA blood velocity to calculated MCA blood flow (panel c). The diagonal line (identity line) indicates where data would fall if there was no measurable effect of protocol on CVR measures (panels a-b) or effect of accounting for MCA CSA in flow calculations (compared to using MCAv alone as an index of flow) when assessing %change of flow or MCAv during SSHC (panel c). Data above the identity line indicates an increase in variable measure with SSHC (compared to ramp; panels a-b) or higher %change in flow value for a given %change in MCAv.

For the steady-state hypercapnia protocol, the difference between the average baseline before start of hypercapnia and the average of the last minute of hypercapnia (i.e., the difference between the SSHC and B conditions in Fig. 4) were calculated. For each individual, the MCA blood velocity CVR during the steady-state hypercapnia condition was then calculated as:

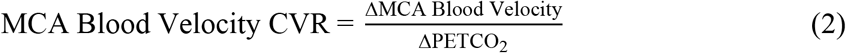

#### MCA blood flow reactivity calculations

The ramp and steady-state hypercapnia MCA blood flow CVR measures are shown in Fig. 5 panel b. For the ramp protocol, the MCA blood velocity was sectioned into 14 second averages corresponding to each CSA image along the ramp protocol. Each of these averaged MCA blood velocity values were multiplied by the corresponding CSA for the given time point to calculate blood flow at 14 second increments along the ramp protocol. To keep it consistent with the MCA blood velocity CVR measures, the PETCO_2_ slopes from the TCD sessions were used as the denominator during CVR calculations. The slopes were calculated for each person and the MCA blood flow CVR was calculated for each individual as:

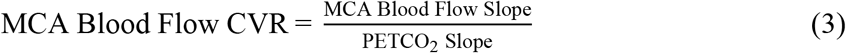

Finally, to compare the MCA blood velocity CVR values and the MCA blood flow CVR values calculated during the steady-state hypercapnia protocol, the percentage change from baseline for each of the MCA blood velocity or blood flow were calculated (Fig. 5, panel c).

### Statistical summary

Inter-rater variability was assessed using Bland-Altman analysis for 45 randomly selected images. Pearson’s correlation coefficient was used to test the correlation for inter-modality (TCD vs. PC-MRI; Fig. 2) MCA blood velocity measures for both baseline and steady-state hypercapnia states (P < 0.05 was considered to be statistically significant using a two-tailed test). In addition, linear slope analysis was conducted to assess changes in PETCO_2_, MAP, MCA CSA, and MCA blood velocity variables during the ramp protocol. A probability level of P < 0.05 indicates a non-zero slope (the linear fits and p-values are labelled on Fig. 3 panels a-e). One-tailed paired t-tests were conducted to compare variable responses during the steady-state hypercapnia condition versus baseline (Fig. 4). Finally, to compare the MCA blood velocity CVR values and the MCA blood flow CVR values calculated during the steady-state hypercapnia protocol, the percentage change from baseline for each of the MCA blood velocity or blood flow was calculated (Fig. 5).

## Results

Our CVR protocols with target PETCO_2_ are illustrated in Fig. 1. Inter-rater variability for CSA measures using the Bland-Altman test indicated a bias of 0.15 ± 0.26 mm^2^ (mean ± S.D.; BKA-MK) and 95% Limits of Agreement from −0.37 to 0.67 mm^2^ (45 randomly selected images). Inter-modality (TCD vs PC-MRI) correlation for MCA blood velocity measures indicated a significant Pearson correlation between baseline and steady-state hypercapnia for each modality (n = 11 participants, 22 pairs; r = 0.69, P < 0.05; Fig. 2).

Slope analysis during the ramp protocol indicated significant slopes (P<0.05; Fig. 3 panels a - d) for achieved ΔPETCO_2_ (mmHg) during TCD (Y = 0.033*X − 3.96; R^2^ = 0.88) and MRI (Y = 0.027*X − 2.54; R^2^ = 0.77) sessions, with corresponding slopes for ΔMCA blood velocity (cm/s; Y = 0.13*X − 13.70; R^2^ = 0.59) and ΔMAP (mmHg; Y = 0.009*X − 0.38; R^2^ = 0.02). Although significant, the slope for MAP during the ramp protocol indicated an average <2mmHg increase throughout the protocol. The slope of the MCA CSA with time did not show a deviation from 0 across the ramp protocol (Fig. 3, panel E; Y = 0.0008*X − 0.057; R^2^ = 0.006).

During the steady-state hypercapnia protocol, PETCO_2_ increased during steady-state hypercapnia from baseline with a mean difference of 4.45 mmHg and 95% CI of 4.05 to 4.85 when using TCD (44.3±3.1 vs. 39.9±2.9 mmHg, respectively; n=12, *η*_p_^2^=0.98, P<0.05; Fig. 4, panel a) and a mean difference of 3.75 and 95% CI of 3.29 to 4.20 when using MRI (42.7±4.2 vs. 38.9±4.4 mmHg, respectively; n=12, *η*_p_^2^=0.97, P<0.05; Fig. 4, panel b) sessions. Similarly, MCA blood velocity increased with steady-state hypercapnia from baseline with a mean difference of 18 cm/s and 95% CI of 13 to 24 cm/s in the TCD trial (104±29 vs. 86±24 cm/s, respectively; n=12, *η*_p_^2^=0.83, P<0.05; Fig. 4, panel c) and a mean difference of 22 cm/s and 95% CI of 12 to 33 when using MRI (via PC-MRI; 79±15 vs. 59±9 cm/s, respectively; n=11, *η*_p_^2^=0.70, P<0.0; Fig. 4 panel e) sessions, and MCA CSA increased with a mean difference of 0.31 mm^2^ and 95% CI of-0.03 to 0.66 mm^2^ during the MRI session (5.71±1.03 vs. 5.34±0.97 mm^2^, respectively; n=12, *η*_p_^2^=0.27, P<0.05; Fig. 4, panel f). The mean difference in MAP between the last minute of steady-state hypercapnia and baseline was 1.35 mmHg with a 95% CI of-0.82 to 3.5 mmHg which is most compatible with a negligible change (90±8 vs. 89±8 mmHg, respectively; n=12; *η*_p_^2^=0.14, Fig. 4, panel e). Calculated MCA blood flows (product of MCA blood velocity and CSA) increased from baseline to steady-state hypercapnia (267±54 vs. 348±78 ml/min, respectively; P<0.05, one-tailed paired t-test) with a delta calculated blood flow of 81±30 ml/min.

The mean difference in MCA blood velocity-based measure of CVR between the ramp and steady-state hypercapnia protocols was 0.18 with 95% CI of −0.59 to 0.96 which is most compatible with a negligible effect of protocol on MCA blood velocity CVR (3.8±1.7 vs. 4.0±1.6 cm/s/mmHg, respectively; n=12, two-tailed paired t-test, p=0.62, *η*_p_^2^=0.02, Fig. 5, panel a). The mean difference in calculated MCA blood flow-based measure of CVR between the steady-state protocol and ramp protocol was 5.0 with a 95% CI of 2.8 to 7.2 ml/min/mmHg and was statistically significant (17.3±5.7 vs. 12.3±4.5 ml/min/mmHg, respectively; n=12, two-tailed t-test, P<0.05, *η*_p_^2^=0.70, two-tailed t-test; Fig. 5, panel b). Similarly, when calculated as %change in MCA blood velocity or MCA blood flow over the ΔPETCO_2_ during hypercapnia, the mean difference between MCA blood flow CVR and MCA blood velocity CVR was (6.7±1.6 vs. 4.8±2.0 %/mmHg, respectively; n = 12, P<.05, one-tailed t-test). Finally, the %change of MCA blood flow was greater than the %change of MCA blood velocity during the steady-state condition with a mean difference of 6.9 and 95% CI of −0.37 to 14.10 (29.8±8.2 vs. 21.5±9.6 %, respectively; n=12, P<0.05, one-tailed t-test; Fig. 5, panel c).

## Discussion

This is the first study to provide MCA CSA measures across ramp and steady-state hypercapnia protocols, enabled by the ability to obtain MCA CSA measurements every 14 seconds with prospective targeting of PETCO_2_. This approach enabled direct comparisons of the steady-state versus a ramp protocols to establish valid CVR calculations using only measures of flow velocity. The noteworthy findings of this study are that: 1) the MCA CSA did not change during the ramp protocol with delta ~ −3 to +4 mmHg of PETCO_2_, but did increase with steady-state hypercapnia ~ +4 mmHg of PETCO_2_), 2) blood pressure remained stable throughout duration of both ramp and steady-state protocols, and 3) CVR measures based on MCA blood velocity cerebrovascular reactivity was not different between ramp and steady-state protocols, but 4) MCA blood flow-based CVR was greater during steady-state compared to the ramp protocol. Taken together, the ramp protocol seems to result in similar CVR values as those observed in the steady-state protocol, with the added advantage of having minimal and negligible effects on MCA CSA or MAP in the face of moderate elevations in PETCO_2_.

### Transcranial Doppler – MCA blood velocity & CVR measures

Often, CVR is characterized by the slope in the linear relationship between MCA blood velocity and PETCO_2_ with respect to each variable’s relative change with time. The relationship between arterial blood velocity and CO_2_ (16) is sigmoidal and both the range and starting point of PETCO_2_ affect the arterial blood velocity response to CO_2_ (17). Regan et al. (7) showed a lower MCA blood velocity-based CVR during steady-state hypercapnia (ΔPETCO_2_ of 10 mmHg) than during the ramp hypercapnic protocol (−5 to +10 mmHg PETCO_2_). However, these ranges of PETCO_2_ can elicit systemic hemodynamic effects (6) that can increase blood velocity changes independent of cerebral vascular bed dilation. The current protocols used a lower dose of change in PETCO_2_ in order to avoid the central hemodynamic and non-linear portions of the CVR curve.

### MRI – MCA CSA & CVR measures

In the current study, PETCO_2_ values at the ends of both ramp and steady-state hypercapnia protocols, and the calculated MCA blood velocity CVR measures, were not different between the two protocols. Yet, the PETCO_2_ changes during the ramp protocol did not affect MCA CSA in same the way that the steady-state hypercapnia protocol did. To quantify transient MCA CSA measures, we developed an anatomical scan optimized to provide MCA images every 14 seconds. This enabled imaging of the MCA during the dynamic stimuli such as the ramp protocol or onset of the steady state protocol, as well as improving temporal sensitivity during the steady-state model. To our knowledge, our 14 second anatomical scan of MCA CSA provides the highest level of temporal resolution (i.e., shortest acquisition time) available when compared to existing assessments of MCA CSA during steady-state hypercapnia protocols. From our current MCA CSA findings, the MCA dilates under conditions where hypercapnia is elevated and sustained (i.e. steady-state hypercapnia), rather than with brief exposure to elevated levels of PETCO_2_ (i.e., end of ramp protocol). This observation is supported by previous studies indicating a slow onset to dilation in either the internal carotid artery (18) or the MCA (2), despite immediate changes in MCA blood velocity that reflect downstream microvascular dilation to early hypercapnia.

This study supports previous MCA CSA findings in CVR studies that indicate MCA CSA dilates after ~2 minutes of steady-state hypercapnia using 3T (2) and 7T MRI (4,5). Thus, CVR values during the steady-state protocol indicated significant error when the MCA cross-sectional area was not included. Specifically, we found that the %change in MCA blood velocity was lower than the %change in MCA blood flow by ~7% (mean difference between these two measures). This value is less than the 18% observed in work by Coverdale et al. (2). We believe the discrepancy in these mean differences is due to the magnitude of hypercapnia achieved in each study, with ~ +Δ4 mmHg PETCO_2_ in the current study versus ~ +Δ10 mmHg in PETCO_2_ for the study by Coverdale et al. As well, there was a 5% increase in MAP in the study by Coverdale et al. (~4 mmHg) which was greater than the increase in MAP for our study (~2 mmHg for ramp, and ~1 mmHg for steady-state), a difference that may be attributed to the greater magnitude of the hypercapnia stimulus in the the previous study. While mild variations in MAP are accounted for by autoregulatory mechanisms such that MCA blood flow is sustained with negligible influence on MCA CSA, autoregulation mechanisms are impaired during hypercapnia (19). Thus, we anticipate that the higher the hypercapnic magnitude, the greater the risk that MAP will influence MCA CSA (and/or blood velocity) augmenting any discrepancies between MCA blood velocity and calculated MCA blood flow.

This is the first study to use a sequential gas delivery circuit (via the RespirAct™) when assessing MCA CSA responses during hypercapnia, thereby more closely aligning partial pressure of arterial and end-tidal CO_2_ levels (20). Interestingly, we achieved similar MCA blood flow CVR during steady-state hypercapnia as previous work (2), and our MCA blood velocity CVR for both the steady-state hypercapnia (2,21) and ramp protocol (7) were in agreement with previous studies. As recommended by Regan et al. (7), when using a limited range of PETCO_2_ values during a hypercapnic protocol, a linear approach to CVR analysis is appropriate.

Everything considered, a CVR measure based solely on TCD-acquired MCA blood velocity measures during a ramp hypercapnia protocol seems to elicit a similar CVR outcome as the commonly used steady-state protocol, but without the limitation of potential changes in MCA CSA. CVR outcome measures involve calculating MCA blood velocity changes for a given change in PETCO_2_. Similarly, our study illustrates that even modest values of change in PETCO_2_ achieve a comparable value of CVR at higher hypercapnic doses.

#### Methodological Considerations

Our overall target range for manipulating PETCO_2_ during the cerebrovascular reactivity was between the −Δ5 to +Δ5 from baseline PETCO_2_ (10). While our findings support a lack of MCA CSA changes during ramp protocols of −Δ3 to +Δ4 mmHg in PETCO_2_, we cannot make conclusive remarks on MCA CSA changes during a ramp protocol of ±Δ5mmHg in PETCO_2_. However, as mentioned above, our values of ΔPETCO_2_ fell within our target ±Δ5mmHg from baseline PETCO_2_ range, the CVR measures are consistent with previous studies where higher levels of PETCO_2_ were used, and we found negligible increases in MAP. Therefore, the modest level of PETCO_2_ achieved here appear to have achieved the major objectives. The ramp protocol designed for the current study also achieved an optimal balance between stimulus and central hemodynamics. We acknowledge however, that we did not account for the impact of the ~2 mmHg rise in MAP during the ramp protocol, which we expect to be negligible in impacting MCA CSA.

Although we were unable to measure continuous blood pressure and MCA blood velocity during the MRI trial, we measured blood velocity data during the MRI session via the PC-MRI sequence. Testing of the TCD and MRI segments of the study were collected consecutively within a 2-hour window with the order of tests varied across participants. The absolute values for MCA blood velocities measured by PC-MRI were lower than our TCD measures, although we suspect this had to do with our pre-set Venc value choice of 100 cm/s which may have cut off some of the higher velocities. Our rationale for not choosing a higher Venc than 100 cm/s was the possibility of cutting off lower velocity values during baseline. Regardless, the increase in MCA blood velocity during hypercapnia (from baseline) were in agreement between the two techniques (TCD: ~Δ18 cm/s and PC-MRI: ~Δ20 cm/s). Thus, we assume that the blood pressure responses during the TCD and MRI sessions were similar as well.

Finally, the current results are delimited to young healthy adults Thus, additional studies are needed to understand the effects of age, a group that demonstrates greater MAP responses to hypercapnia and variable responses in CSA changes (9), or other differentiating conditions. Further, our study was conducted in the supine position, which is less replicable for CVR studies than the seated position (22) and may explain some of the disparity of blood velocity measures for some individuals between the two protocols: the supine posture is necessary for MRI studies. Additional studies are required to address the impact of CSA changes and protocol model on posture-dependent intra-subject variability so that reliable CVR protocols can be used to make inferences on vascular health (e.g., inferred vascular dysfunction with reduced CVR). The current data suggest that the ramp protocol might be useful in reducing inter-study variability.

## Conclusions

A constant CSA during experimental vasoreactive challenges is essential to the reliability of TCD-acquired blood velocity as a correlative index of blood flow changes. In this study, we showed data that were most compatible with negligible change in MCA CSA from baseline during a graded ramp hypercapnic protocol (± ~Δ4 mmHg from baseline PETCO_2_). Similar to our previous work, the MCA data align best with an interpretation of MCA dilation during a prolonged (3-minutes) ~Δ4 mmHg in baseline PETCO_2_ period of hypercapnia, and confirmed the expected error in %change of MCA blood velocity as an index of blood flow and in the calculated CVR when a change in CSA is not considered (2). Combined, these data suggest that during transient changes in PETCO_2_, as in the ramp protocol, the constant segment of the sigmoid may describe MCA CSA changes with PETCO_2_, and any changes in blood velocity can reflect CBF. In summary, the ramp protocol with ± ~Δ4 mmHg from baseline PETCO_2_ appears to provide expected measures of CVR where blood velocity should reflect blood flow patterns of change.

## Acknowledgements

The authors would like to thank the participants for their time, Joseph S. Gati and Trevor Szekeres for their MRI expertise, Arlene Fleischhauer, and Emilie Woehrle and Jenna Schulz for helping with participant recruitment. This study was funded by the Canadian Institutes of Health Research (grant # 201503MOP-342412-MOV-CEEA). KN was supported by a research grant from Children’s Health Foundation. BKA was funded by the MITACS Postdoctoral Elevate Fellowship. JKS and RSM are Tier 1 Canada Research Chairs.

## Author contribution statement

BKA conceptualized and designed the study, collected, analyzed, and disseminated the data, and wrote and edited the manuscript. SB assisted with study design, data collection and analysis, and
assisted with writing and editing the manuscript. MK assisted with data collection, analysis and interpretation and editing of the manuscript. BJM assisted with data collection, organization, and editing the manuscript. KN assisted with study design and editing of the manuscript. RSM assisted with study design and editing of the manuscript. JKS conceptualized and designed the study, and assisted with data dissemination, writing and editing of the manuscript.

## Conflict of interest

The authors do not have any conflicts of interest to disclose.

## Notes

### Competing Interest Statement

The authors have declared no competing interest.

## References

1. Portegies MLP, de Bruijn RFAG, Hofman A, Koudstaal PJ, Ikram MA. Cerebral vasomotor reactivity and risk of mortality: the Rotterdam Study. Stroke. 2014 Jan; 45(1):42–7.

2. Coverdale NS, Gati JS, Opalevych O, Perrotta A, Shoemaker JK. Cerebral blood flow velocity underestimates cerebral blood flow during modest hypercapnia and hypocapnia. J Appl Physiol Bethesda Md 1985. 2014 Nov 15; 117(10):1090–6.

3. Howe CA, Caldwell HG, Carr J, Nowak-Flück D, Ainslie PN, Hoiland RL. Cerebrovascular reactivity to carbon dioxide is not influenced by variability in the ventilatory sensitivity to carbon dioxide. Exp Physiol. 2020; 105(5):904–15.

4. Al-Khazraji BK, Shoemaker LN, Gati JS, Szekeres T, Shoemaker JK. Reactivity of larger intracranial arteries using 7 T MRI in young adults. J Cereb Blood Flow Metab Off J Int Soc Cereb Blood Flow Metab. 2019 Jul; 39(7):1204–14.

5. Verbree J, Bronzwaer A-SGT, Ghariq E, Versluis MJ, Daemen MJAP, van Buchem MA, et al. Assessment of middle cerebral artery diameter during hypocapnia and hypercapnia in humans using ultra-high-field MRI. J Appl Physiol Bethesda Md 1985. 2014 Nov 15; 117(10):1084–9.

6. Shoemaker JK, Vovk A, Cunningham DA. Peripheral chemoreceptor contributions to sympathetic and cardiovascular responses during hypercapnia. Can J Physiol Pharmacol. 2002 Dec; 80(12):1136–44.

7. Regan RE, Fisher JA, Duffin J. Factors affecting the determination of cerebrovascular reactivity. Brain Behav. 2014 Sep; 4(5):775–88.

8. Stefanidis KB, Askew CD, Klein T, Lagopoulos J, Summers MJ. Healthy aging affects cerebrovascular reactivity and pressure-flow responses, but not neurovascular coupling: A cross-sectional study. PLOS ONE. 2019 May 16; 14(5):e0217082.

9. Coverdale NS, Badrov MB, Shoemaker JK. Impact of age on cerebrovascular dilation versus reactivity to hypercapnia. J Cereb Blood Flow Metab. 2017 Jan 1; 37(1):344–55.

10. Hoiland RL, Fisher JA, Ainslie PN. Regulation of the Cerebral Circulation by Arterial Carbon Dioxide. Compr Physiol. 2019 Jun 12; 9(3):1101–54.

11. McKetton L, Cohn M, Tang-Wai DF, Sobczyk O, Duffin J, Holmes KR, et al. Cerebrovascular Resistance in Healthy Aging and Mild Cognitive Impairment. Front Aging Neurosci. 2019;11:79.

12. Fisher JA, Sobczyk O, Crawley A, Poublanc J, Dufort P, Venkatraghavan L, et al. Assessing cerebrovascular reactivity by the pattern of response to progressive hypercapnia. Hum Brain Mapp. 2017 Jul; 38(7):3415–27.

13. Brothers RM, Lucas RAI, Zhu Y-S, Crandall CG, Zhang R. Cerebral vasomotor reactivity: steady-state versus transient changes in carbon dioxide tension. Exp Physiol. 2014 Nov; 99(11):1499–510.

14. Core Team R. R: A language and environment for statistical computing. R Foundation for Statistical Computing, Vienna, Austria (2013). Suppl Fig S. 2015;2.

15. Makiyama K. magicfor: Magic Functions to Obtain Results from for Loops [Internet]. 2016 [cited 2021 Jan 9]. Available from: https://CRAN.R-project.org/package=magicfor

16. Battisti-Charbonney A, Fisher J, Duffin J. The cerebrovascular response to carbon dioxide in humans. J Physiol. 2011 Jun 15; 589(Pt 12):3039–48.

17. Sobczyk O, Battisti-Charbonney A, Fierstra J, Mandell DM, Poublanc J, Crawley AP, et al. A conceptual model for CO_2_-induced redistribution of cerebral blood flow with experimental confirmation using BOLD MRI. NeuroImage. 2014 May 15; 92:56–68.

18. Willie CK, Macleod DB, Shaw AD, Smith KJ, Tzeng YC, Eves ND, et al. Regional brain blood flow in man during acute changes in arterial blood gases. J Physiol. 2012 Jul 15; 590(14):3261–75.

19. Panerai RB, Deverson ST, Mahony P, Hayes P, Evans DH. Effects of CO2 on dynamic cerebral autoregulation measurement. Physiol Meas. 1999 Aug; 20(3):265–75.

20. Ito S, Mardimae A, Han J, Duffin J, Wells G, Fedorko L, et al. Non-invasive prospective targeting of arterial P(CO2) in subjects at rest. J Physiol. 2008 Aug 1; 586(15):3675–82.

21. Burley CV, Lucas RAI, Whittaker AC, Mullinger K, Lucas SJE. The CO2 stimulus duration and steady-state time point used for data extraction alters the cerebrovascular reactivity outcome measure. Exp Physiol. 2020 May; 105(5):893–903.

22. McDonnell MN, Berry NM, Cutting MA, Keage HA, Buckley JD, Howe PRC. Transcranial Doppler ultrasound to assess cerebrovascular reactivity: reliability, reproducibility and effect of posture. PeerJ. 2013 Apr 9; 1:e65.

